# Altered effective connectivity from the posterior insula to the amygdala mediates the relationship between psychopathic traits and concern with the Harm foundation

**DOI:** 10.1101/2021.06.21.449236

**Authors:** Shuer Ye, Wei Li, Bing Zhu, Yating Lv, Qun Yang, Frank Krueger

**Author notes:** **Corresponding authors:** Yating Lv,; Qun Yang. The two authors contributed equally to the research (Shuer Ye, Wei Li).

## Abstract

Psychopathic traits have been demonstrated to be associated with different types of morality; however, the neuropsychological mechanism underlying the relationship between psychopathic traits and morality remains unclear. Our study examined the effective connectivity (EC) of psychopathic traits-related brain regions and its association to concern with different moral foundations by combining behavioral measures with resting-state fMRI. We administered the Levenson Self-Report Psychopathy Scale (LSRP) and Moral Foundation Questionnaire (MFQ) to 78 college students after resting-state fMRI scanning. Our results showed that total and primary psychopathic traits score predicted concern with the Harm foundation. The EC from the posterior insula to the amygdala was negatively correlated with psychopathic traits and positively with concern with the Harm foundation. Altered posterior insula-amygdala EC partially mediated the relationship between psychopathic traits and concern with the Harm foundation. Our findings indicated that individuals with elevated psychopathic traits may have atypical processes in recognizing and integrating bodily state information into emotional responses, leading to less concern for harm-related morality. The study deepened our understanding of the neuropsychological mechanism underlying the relationship between psychopathic traits and morality and may have implications for the prevention of higher psychopathic traits individuals from committing serious transgressions.

## INTRODUCTION

Psychopathy has been recognized as a personality disorder, primarily characterized by problematic interpersonal relations, shallow affect, poor behavioral control, irresponsibility, and disregard of moral norms (Hare, 1991, 2003). There are two subtypes of psychopathy, including primary psychopathy and secondary psychopathy. Primary psychopathy is associated with interpersonal and affective features whereas secondary psychopathy is linked to aggressive and antisocial lifestyles (Hare, 1991). Psychopathy has been increasingly recognized as a continuum of personality traits rather than a taxonomic feature (Corr, 2010). Those who are not clinically identified as psychopaths, but with higher psychopathic traits, may also show typical psychopathic characteristics (Brennan et al., 2018; Djeriouat & Trémolière, 2014; Neumann & Hare, 2008).

Individuals with elevated psychopathic traits may engage in various immoral behaviors, posing challenges to societies in maintaining social order (Beaver et al., 2017). Therefore, the relationship between psychopathic traits and moral cognition is of great interest to many researchers (Blair, 2007; Gao & Tang, 2013; Marsh et al., 2011). Moral Foundation Theory (MFT), a widely known framework on human morality, provides a new perspective for investigating the relationship between psychopathic traits and morality (Gay et al., 2018). According to this theory, morality consists of five foundations: (1) Harm (preventing harm to others); (2) Fairness (preserving fairness, equal rights, and justice); (3) Loyalty (practicing loyalty to one’s own group); (4) Authority (respecting authority and social order); and (5) Purity (pursuing purity or sanctity of body, mind, and soul) (Haidt, 2012; Haidt & Graham, 2007).

Psychopathic traits have been consistently shown to be associated with concern with different moral foundations (Aharoni et al., 2011; Fernandes et al., 2020). This relation was especially pronounced for the Harm and Fairness foundations (Aharoni et al., 2011; Glenn et al., 2009a; Marshall et al., 2018). Individuals scoring higher in psychopathic traits are significantly less likely to endorse the Harm and Fairness foundations in the community (Glenn et al., 2009a) and forensic samples (Aharoni et al., 2011). Furthermore, emotion may serve as a mediator in the relationship between psychopathic traits and morality (Glenn et al., 2009a; Patil, 2015; Ye et al., 2021). For example, a lower level of empathic concern predicts less endorsement for preventing harm and unfairness in individuals with elevated psychopathic traits (Glenn et al., 2009a). Our recent study showed that unpleasantness plays a mediating role in psychopathic traits and moral judgment especially for the Harm and Fairness foundations (Ye et al., 2021). However, few studies have yet directly examined the underlying neural substrates of the relationship between psychopathic traits and concern with the moral foundations. Our current paper aims to explore the neural mechanism underlying the relationship between psychopathic traits and morality under the framework of MFT.

To date, much of our knowledge about the neural mechanism of the relationship between psychopathic traits and morality is from task-dependent fMRI studies, which associate blood oxygenation level-dependent (BOLD) signals with moral judgments for individuals with higher psychopathic traits. In particular, many of these studies revealed altered brain functions in brain regions supporting emotion processing, such as the amygdala. For example, individuals with elevated psychopathic traits show decreased activation in the amygdala during moral decision-making (Glenn et al., 2009b; Harenskia et al., 2014; Yoder et al., 2015). Youths with higher psychopathic traits show reduced functional connectivity between the amygdala and orbitofrontal cortex (OFC) when making moral judgments (Marsh et al., 2011). In addition, male inmates with higher psychopathic traits demonstrate decreased functional connectivity between the amygdala and right temporoparietal junction when viewing morally bad scenarios (Yoder et al., 2015).

Task-independent resting-state fMRI (RS-fMRI) offers an appealing alternative to characterize the neural functions of individuals with mental health concerns (Philippi et al., 2015). RS-fMRI measures intrinsic brain functions by scanning participants while they maintain a relatively relaxed state in a scanner. Compared with task-fMRI, RS-fMRI measures have a better signal to noise ratio and have good reliability (Fox & Greicius, 2010). Recent evidence has shown altered resting-state architecture of functional brain organization associated with psychopathic traits. In particular, abnormal functional connectivity between limbic-paralimbic structures (e.g., insula, amygdala, and posterior cingulate cortex [PCC]) and prefrontal areas (e.g., ventromedial prefrontal cortex [vmPFC], dorsal medial prefrontal cortex [dmPFC], ventrolateral prefrontal cortex, and dorsolateral prefrontal cortex) have been shown to be related to elevated psychopathic traits (Contreras-Rodriguez et al., 2014; Motzkin et al., 2011). Overall, RS-fMRI findings also reveal altered functional connectivity between emotional-related brain regions (e.g., the amygdala) and other brain areas including the prefrontal areas for individuals with higher psychopathic traits (Blair, 2007; Contreras-Rodriguez et al., 2014; Espinoza et al., 2019; Motzkin et al., 2011). However, those RS-fMRI studies have provided little information about how resting-state functional connectivity between those regions of interest underlie the relationship of psychopathic traits to moral cognition. Especially, little is known about the direction of these connections (i.e., functional connectivity, FC) during moral cognition for people with elevated psychopathic traits. Finding out the direction of the information transmission between brain regions might deepen our understanding of the neural mechanism underlying the moral cognition for people with elevated psychopathic traits. For example, given the atypical emotional processing widely associated with psychopathic traits, we might want to know whether it is because brain regions supporting emotional functions send or receive atypical inputs from other brain regions.

Effective connectivity (EC) provides an estimation of the directional relationship between brain regions. Therefore, information about the possible causal influence of one region on another can be inferred (Friston & Moran, 2013). Granger causality analysis (GCA) is a data-driven method for investigating the effective connectivity between recorded time series of brain regions without a prior specification of causal setting (Seth et al., 2015). Using this method might increase our knowledge about the atypical neural circuits associated with moral cognition for individuals with higher psychopathic traits in terms of causal relationships between brain regions without a prior hypothesis of the brain connectivity models.

In our study, we aimed to utilize a GCA approach to explore the intrinsic neural signatures for the relationship between psychopathic traits and concern with different moral foundations. We focused on brain regions that have been proved to be significantly associated with psychopathic traits in a recent meta-analysis including the amygdala, dmPFC, PCC, anterior cingulate cortex (ACC), OFC, and hippocampus (Deming & Koenigs, 2020). Given the essential role of emotion in the relationship between psychopathic traits and moral cognition (Glenn et al., 2009a; Patil, 2015; Ye et al., 2021), we expected that the EC from brain regions that are particularly related to emotional function (e.g., amygdala) to other brain regions might mediate the relationship between psychopathic traits and moral foundations, especially for the Harm and Fairness foundations. In other words, those brain areas supporting emotional function might exert atypical influence on other brain regions during moral cognition for individuals with elevated psychopathic traits. Additionally, We expected our findings to be more pronounced for primary psychopathy than secondary psychopathy as recently we found psychopathic traits are correlated with moral judgments merely in primary psychopathy (Ye et al., 2021).

## METHOD

### Participants

Seventy-eight college students from local universities participated in this study. All the participants were right-handed and in healthy condition. No participants reported any history of neurological or psychiatric disorder. During the study, one participant failed to complete the questionnaires, five participants were detected to provide inappropriate answers in the bogus items in MFQ, and two participants had head motions exceeding 2° during scanning. These participants were excluded from further analyses, leaving a final sample of 70 participants (29 males; 22.71 ± 2.51 years old, range: 18–30 years old) (**Tab.1**). The study was approved by the Ethical Committee of Hangzhou Normal University. All the participants provided written informed consent in accordance with the Declaration of Helsinki and were compensated by cash.

**Table 1.** Demographic information.

|  | Male (N=29) |  | Female (N=41) |  | *p*-value |
| --- | --- | --- | --- | --- | --- |
|  | Mean | SD | Mean | SD |  |
| Age (years) | 23.52 | 2.86 | 22.15 | 2.08 |  |
| T-PTS | 52.34 | 7.41 | 53.68 | 7.70 | 0.469 |
| P-PTS | 32.69 | 5.57 | 33.00 | 5.10 | 0.810 |
| S-PTS | 19.66 | 3.15 | 20.68 | 3.56 | 0.217 |
| Harm foundation | 19.10 | 3.93 | 20.49 | 3.24 | 0.112 |
| Fairness foundation | 19.72 | 4.57 | 20.56 | 3.10 | 0.363 |
| Loyalty foundation | 18.24 | 3.47 | 20.17 | 3.71 | 0.031<sup>*</sup> |
| Authority foundation | 16.21 | 4.72 | 17.68 | 3.54 | 0.139 |
| Purity foundation | 15.38 | 4.59 | 16.63 | 4.02 | 0.229 |

SD: standard deviation; LSRP: the Levenson Self-Report Psychopathy Scale; T-TPS: total psychopathic traits score; P-PTS: primary psychopathic traits score; S-PTS: secondary psychopathic traits score.
\*: $p < 0.05$

### Measures and Procedure

#### Measures

##### Levenson Self-Report Psychopathy Scale (LSRP)

Psychopathic traits were assessed with the LSRP which has been well validated in the community sample (Levenson et al., 1995). The scale consists of 26 items, each of which can be scored on a 4-point Likert scale ranging from 1 (strongly disagree) to 4 (strongly agree). The questionnaire includes two separate subtypes of psychopathy (i.e., primary psychopathy and secondary psychopathy). Primary psychopathy corresponds to Factor 1 (interpersonal/affective traits) whereas secondary psychopathy corresponds to Factor 2 (lifestyle/antisocial traits) of The Psychopathy Checklist-Revised which is a structured clinical assessment. The higher scores on the scale indicate higher psychopathic traits (Lynam et al., 1999). The Chinese version of the LSRP has been proved to have good reliability (M.-C. Wang et al., 2018). The Cronbach’s alpha was 0.76 (total psychopathic traits score [T-PTS]), 0.72 (primary psychopathic traits score [P-PTS]) and 0.59 (secondary psychopathic traits score [S-PTS]) in the present sample.

#### Moral Foundation Questionnaire (MFQ)

The concern with different moral foundations were evaluated by the MFQ, which consists of 32 items (Graham et al., 2009). The scale contains two sections. The first section measures the moral relevance of different moral considerations on a six-point Likert scale ranging from 0 (not at all relevant) to 5 (extremely relevant) while the second section measures the degree to which participants agreed or disagreed with stated moral views on a six-point Likert scale ranging from 0 (strongly disagree) to 5 (strongly agree). The scores of five moral foundations (i.e., Harm, Fairness, Loyalty, Authority, and Purity) were calculated respectively, with higher scores indicating higher concern of the corresponding moral foundation. The questionnaire has been well-validated in Chinese sample (Du, 2019; R. Wang et al., 2019). The Cronbach’s alpha of subscales was 0.41 (Harm), 0.58 (Fairness), 0.53 (Loyalty), 0.61(Authority) and 0.62 (Purity) in the present sample.

#### Procedure

Participants completed two questionnaires after undergoing task-free fMRI. The order of the two questionnaires were counterbalanced among participants.

### Behavioral data analyses

The statistical analyses for questionnaire measures were performed using SPSS Version 23.0 (IBM Corp. Released 2015) with an alpha value of p < 0.05 (two tailed). Linear regression analyses were performed to examine whether total LSRP score, primary psychopathy, and secondary psychopathy could predict concern with the five moral foundations after controlling the demographic regressors (i.e., age and gender).

### Imaging acquisition

All MRI data were acquired at the Center for Cognition and Brain Disorders of Hangzhou Normal University using a 3T MRI scanner (GE Discovery 750 MRI, General Electric, Milwaukee, WI, USA). The participants were scanned for 8 minutes and instructed to keep their eyes closed and remain awake without performing any systematic thinking. Head motion was minimized using foam padding and restraint. The imaging parameters of the EPI sequence were as follows: repetition time (TR) = 2000ms; interleaved 43 slices; echo time (TE) = 30 ms; thickness = 3.2mm; flip angle = 90°; field of view (FOV) = 220 ×220 mm2; and matrix size= 64 × 64. Each fMRI scan included 240 imaging volumes. In addition, a high-resolution 3D T1-weighted anatomical image was also acquired using a magnetization prepared gradient echo (MP-RAGE) sequence with following imaging parameters: 176 sagittal slices; TR = 8,100ms; slice thickness = 1 mm; TE = 3.1ms; flip angle = 8°; FOV = 250 × 250 mm.

### Preprocessing

The neuroimaging data were preprocessed using the DPABI software package based on SPM12 (Yan et al., 2016). The first 10 volumes of the functional images were discarded to eliminate the non-equilibrium effects of magnetization. Then, slice timing and head motion correction were applied. Corrected functional images were registered to the corresponding T1-weighted anatomical image and spatially transformed to the standard MNI space with 3 × 3 × 3 mm3 voxels. A band-pass temporal filtering (0.01–0.1 Hz) was subsequently applied to the time series to reduce the effect of low-frequency drifts and high-frequency physiological noise. Next, the images were spatially smoothed using an isotropic Gaussian kernel with 4mm full width at half maximum to decrease spatial noise. Finally, the linear trends of time courses were removed, and three common nuisance variables including 24 head-motion parameters and averaged signals from the cerebrospinal fluid and white matter were regressed out.

### Selection of the regions of interest

To access the EC of the regions that are most relevant to psychopathic traits, a priori seed regions of interest (ROIs) were selected that have been identified from a meta-analysis (Deming & Koenigs, 2020): anterior cingulate cortex (ACC), dorsomedial prefrontal cortex (dmPFC), inferior frontal gyrus (IFG), posterior orbitofrontal cortex (OFC), amygdala, hippocampus, and calcarine (**Tab.2**). Each ROI was centered on the coordinates in MNI space provided by the meta-analysis and include a radius sphere of 5 mm (**Fig.1b**).

**Table 2.** Seeds selected to evaluate effective connectivity.

| Seed regions | Hemi. | MNI coordinates |  |  |
| --- | --- | --- | --- | --- |
|  |  | x | y | z |
| Anterior cingulate cortex | L | -6 | 32 | 22 |
| Dorsomedial prefrontal cortex 1 | L/R | -2 | 56 | 28 |
| Dorsomedial prefrontal cortex 2 | L/R | -2 | 50 | 34 |
| Dorsomedial prefrontal cortex 3 | L/R | 2 | 44 | 32 |
| Inferior frontal gyrus | R | 40 | 40 | -8 |
| Posterior orbitofrontal cortex | R | 26 | 8 | -16 |
| Amygdala | R | 30 | 4 | -20 |
| Hippocampus 1 | R | 24 | -36 | 2 |
| Hippocampus 2 | R | 30 | -40 | -4 |
| Calcarine | R | 12 | -70 | 16 |

L: left; R: right

**Figure 1.**
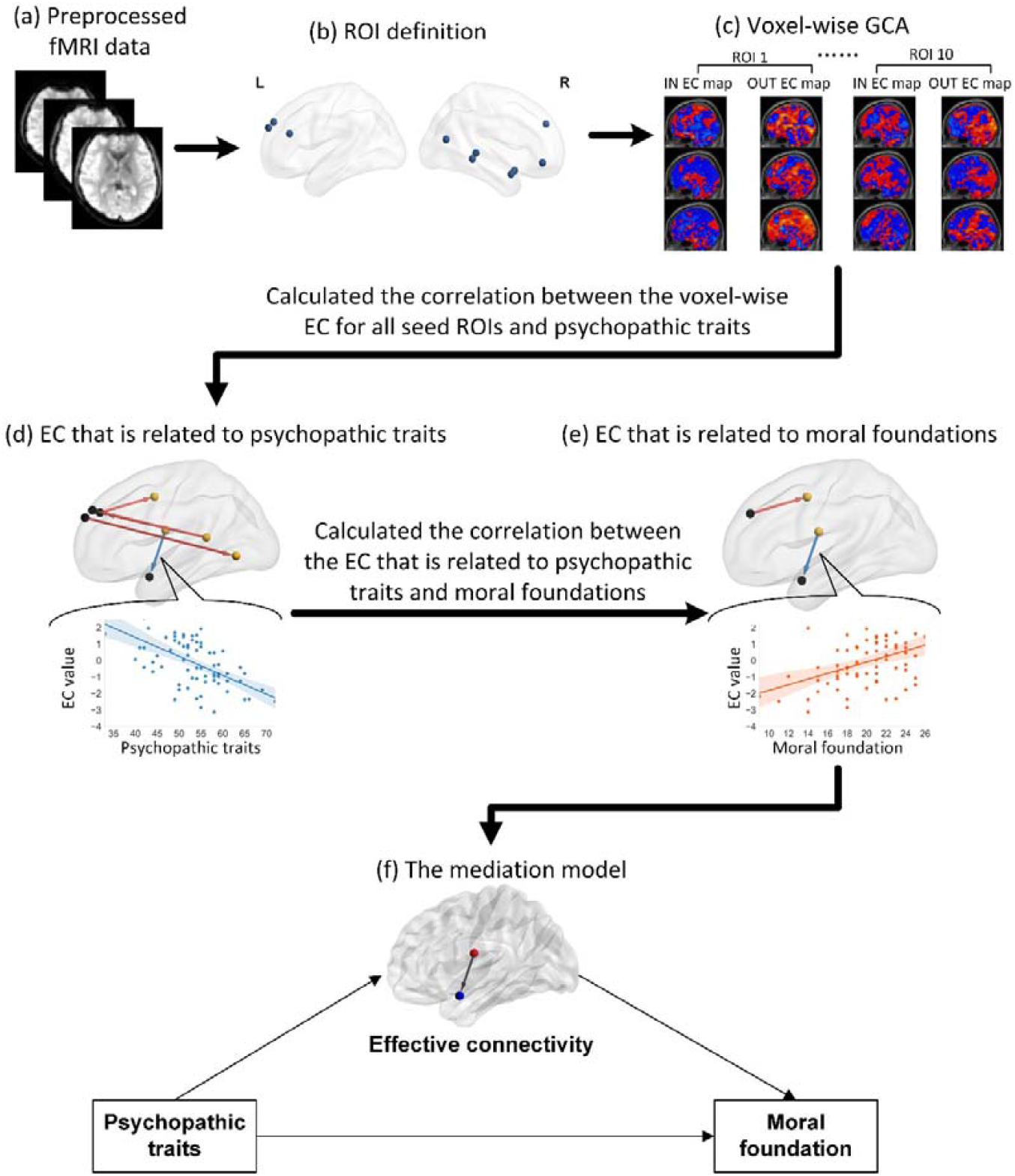
Workflow of the neuroimaging data analyses. First, preprocessed rs-fMRI data were used to calculate voxel-wise EC based on ten predefined ROIs (subplot a, b, and c). Then, partial correlation analyses were performed between EC and scores on psychopathic traits (i.e. T-PTS, P-PTS, and S-PTS) (subplot d). Next, partial correlation analyses were performed between EC that is significantly related to psychopathic traits and concerns with moral foundations (subplot e). Finally, mediation analyses were conducted to examine the mediation role of these EC in the relationship between psychopathic traits and concern with moral foundations (subplot f). T-PTS: total psychopathic traits score; P-PTS: primary psychopathic traits score; S-PTS: secondary psychopathic traits score.

### Granger causality analysis

Bivariate coefficient-based GCAs were performed to examine EC among ROIs and each voxel of the whole brain using the REST v1.8 toolbox (Song et al., 2011; Zang et al., 2012). This GCA approach assumes that if including past values of a BOLD time series x lead to better prediction of the current value of time series y, then series x causes series y (Roebroeck et al., 2005). Here, the BOLD time series of each ROI was defined as the seed time series x, and the time series of voxels within the whole brain were defined as the time series y. The positive value of EC from series x to series y indicates an excitatory effect that series x imposes on series y, whereas the negative value was interpreted as an inhibitory effect (Zang et al., 2012). Finally, Fisher’s r-to-z transformations were implemented (**Fig.1c**).

### Partial correlation analyses

Voxel-wise EC for all seed ROIs significantly associated with psychopathic traits (T-PTS, P-PTS, and S-PTS) were examined by using partial correlation analyses with age and gender as covariates (**Fig.1d**). To correct for multiple comparisons, family-wise error (FWE) correction with Gaussian random field theory was applied, in which the voxel threshold was set to p < 0.001 and the cluster threshold was set to p < 0.05 with two tails. Next, the relationship between the neuroimaging findings (i.e., EC significantly related to psychopathic traits) and concern with the five moral foundations was examined (**Fig.1e**). The values of EC that was significantly related to psychopathic traits were extracted. Then, Pearson correlation analyses were further conducted to examine the relationship between these EC values and concern with the five moral foundations with age and gender as covariates. Here, Bonferroni correction for multiple comparisons (n = 65) was applied to control the false-positive results.

### Mediation analyses

For the EC that was significantly associated with both psychopathic traits and the concern with the five foundations, mediation analyses were conducted to examine whether relationship between psychopathic traits and moral concern was mediated by the EC (**Fig.1f**). The mediation analyses were conducted by applying Bootstrapping PROCESS for SPSS with age and gender as covariates. The Bootstrap samples were set as 5,000, and the confidence level (CI) for confidence intervals was set as 95%. The CI which does not include zero suggests a significant mediation role in the relationship between psychopathic traits and the concern with the five foundations.

## RESULTS

### Relationship between psychopathic traits and five moral foundations

The relationship between psychopathic traits and concerns with the moral foundations were examined by establishing linear regression models. The results showed that concern with the Harm foundation was significantly predicted by T-PTS (β = -0.461, *p* < 0.001) and P-PTS (β = -0.498, *p* < 0.001). However, psychopathic traits did not predict concern with the other four foundations (i.e., Fairness, Loyalty, Authority and Purity) (**Tab. 3**).

**Table 3.** Regression coefficients for the T-PTS, P-PTS, S-PTS, and concern with five moral foundations.

|  | Regression coefficient |  |  |
| --- | --- | --- | --- |
|  | T-PTS | P-PTS | S-PTS |
| Harm | -0.461*** | -0.498*** | -0.258 |
| Fairness | -0.174 | -0.243 | -0.011 |
| Loyalty | -0.072 | -0.036 | -0.107 |
| Authority | -0.066 | -0.079 | -0.024 |
| Purity | -0.226 | -0.156 | -0.266 |

\*\*\* Bonferroni-corrected $p < 0.001$ , $n=15$ ; T-PTS: total psychopathic traits score; P-PTS: primary psychopathic traits score; S-PTS: secondary psychopathic traits score.

### Relationship between psychopathic traits and EC

Partial correlation analyses were conducted to examine the voxel-wise EC that were significantly associated with psychopathic traits. The results showed that T-PTS was significantly related to the EC from the posterior insula to the amygdala, from the dmPFC to the middle temporal gyrus and precentral gyrus, and from the PCC to the dmPFC (**Fig.2**). P-PTS was significantly correlated with the EC from the posterior insula and supramarginal gyrus to the amygdala, from the dmPFC to the precentral gyrus and IPL, from the PCC to the dmPFC, and from the ACC to the middle temporal gyrus **(Fig.3)**. However, no EC was found to be significant correlated with S-PTS. The detailed results for the relationship between the EC and psychopathic traits are shown in Table 4.

**Figure 2.**
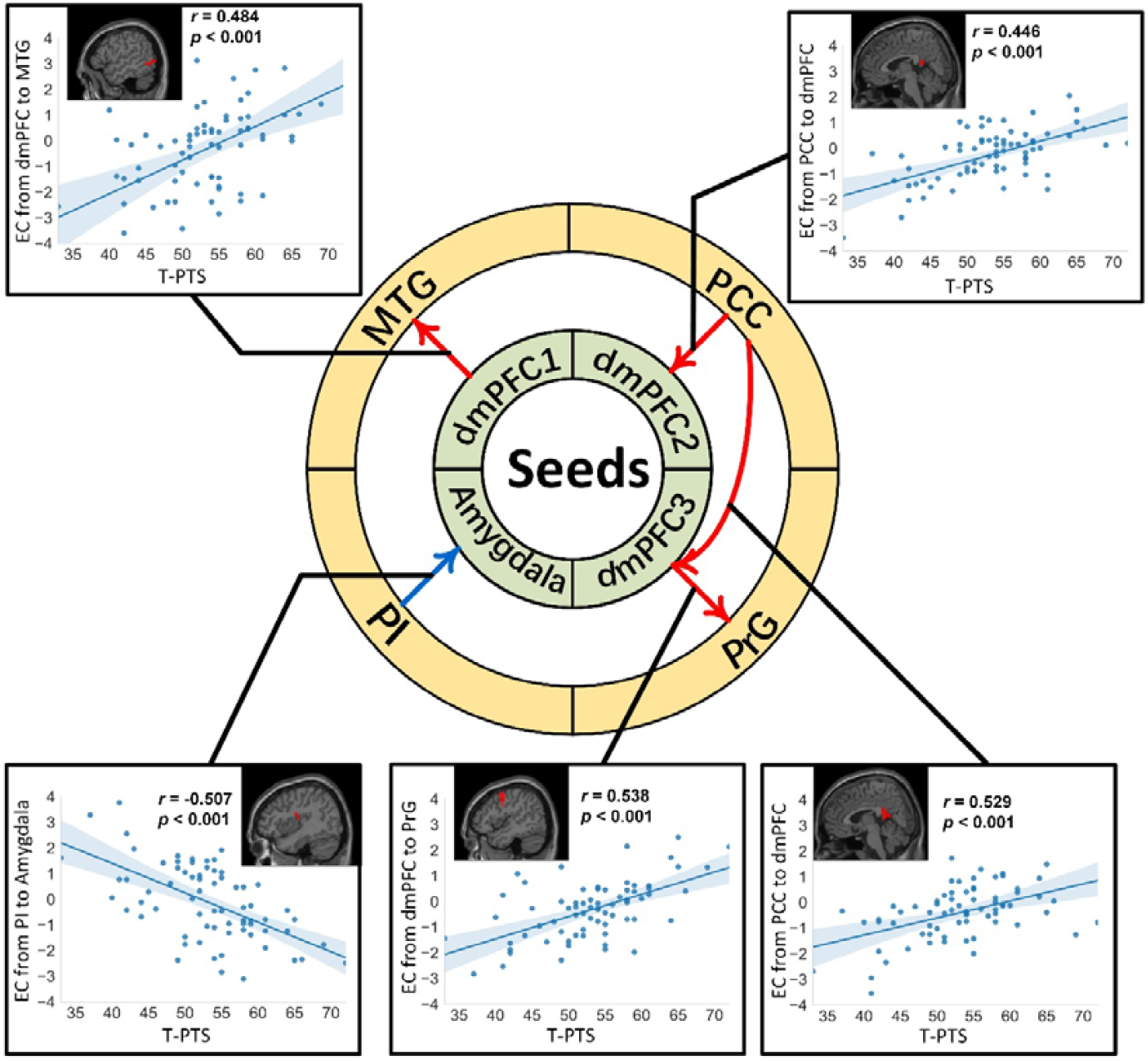
EC that is significantly related to T-PTS. By conducting partial correlation analyses, the EC from the dmPFC to the MTG and PrG, from the PCC to the dmPFC, and from the PI to the amygdala were found to be significantly related to T-PTS. The red arrow refers to a positive association between EC and T-PTS while the blue arrow refers to a negative association between EC and T-PTS. EC: effective connectivity; MTG: middle temporal gyrus; PCC: posterior cingulate cortex; PrG: precentral gyrus; PI: posterior insula; dmPFC: dorsomedial prefrontal cortex; T-PTS: total psychopathic traits score.

**Figure 3.**
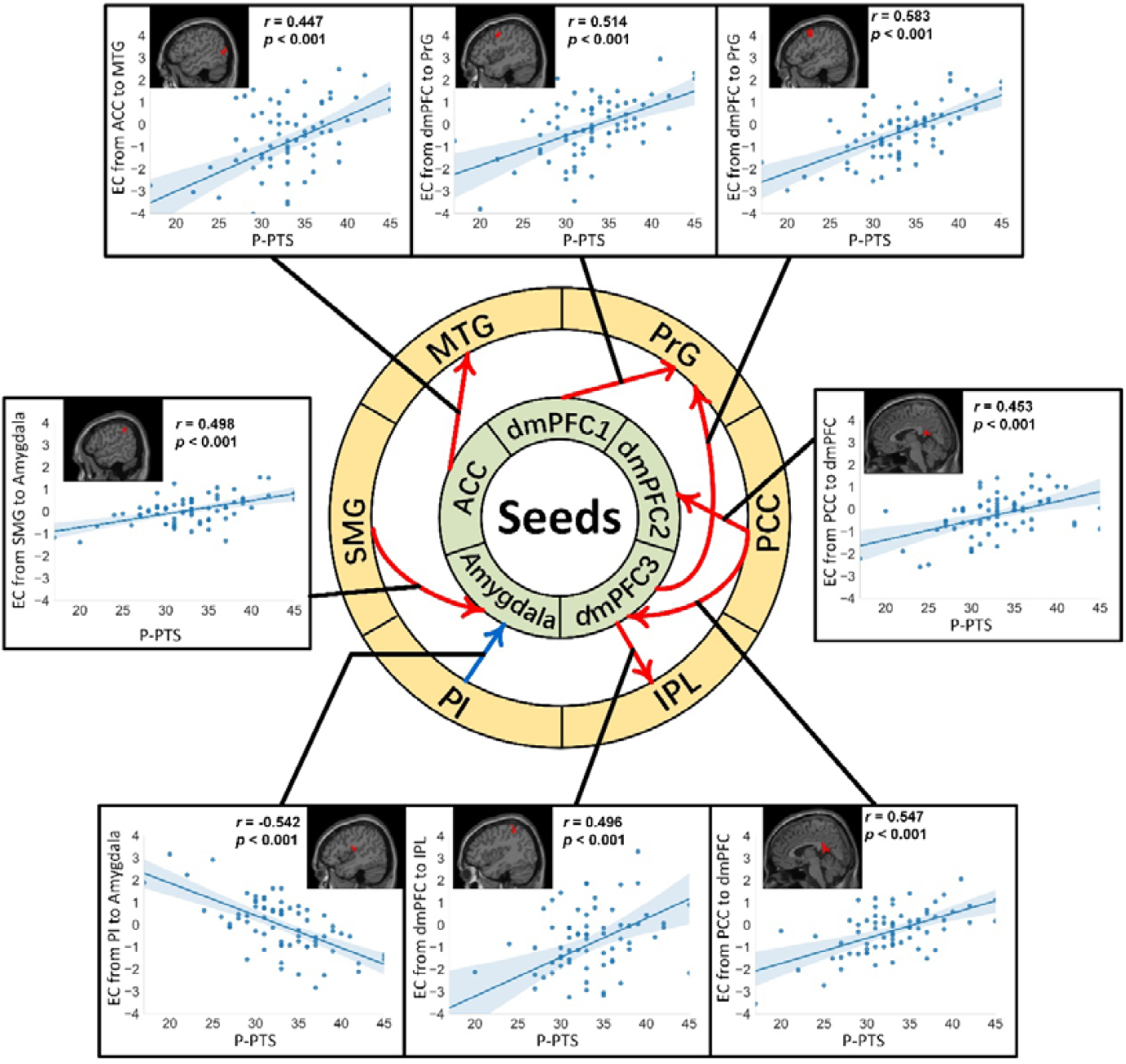
EC that is significantly related to P-PTS. By conducting partial correlation analyses, EC from the ACC to the MTG, form the dmPFC to the PrG and IPL, from the PCC to the dmPFC, and from the PI and SMG to the amygdala were found to be significantly related to P-PTS. The red arrow refers to a positive association between EC and P-PTS while the blue arrow refers to a negative association between EC and P-PTS. EC: effective connectivity; MTG: middle temporal gyrus; PCC: posterior cingulate cortex; PrG: precentral gyrus; PI: posterior insula; IPL: inferior parietal lobe; SMG: supramarginal cortex; ACC: anterior cingulate cortex; dmPFC: dorsomedial prefrontal cortex; P-PTS: primary psychopathic traits score.

**Table 4.** The Effective connectivity associated with psychopathic traits and their relationship to moral concern with moral foundations.

| Seed regions | Direction | Brain regions | MNI coordinate |  |  | Peak Intensity | Cluster Size <sup>a</sup> | *r*-value |  |  |  |  |
| --- | --- | --- | --- | --- | --- | --- | --- | --- | --- | --- | --- | --- |
|  |  |  | x | y | z |  |  | Harm | Fairness | Loyalty | Authority | Sanctity |
| ***T-PTS*** |  |  |  |  |  |  |  |  |  |  |  |  |
| Dorsomedial prefrontal cortex 1 | Output | Middle temporal gyrus | -57 | -66 | -3 | 0.484 | 47 | -0.130 | -0.093 | 0.132 | -0.013 | -0.117 |
| Dorsomedial prefrontal cortex 2 | Input | Posterior cingulate cortex | -3 | -42 | 24 | 0.446 | 28 | -0.260 | -0.277 | 0.158 | 0.051 | -0.011 |
| Dorsomedial prefrontal cortex 3 | Input | Posterior cingulate cortex | 6 | -42 | 12 | 0.529 | 144 | -0.248 | -0.063 | 0.057 | -0.037 | -0.154 |
| Dorsomedial prefrontal cortex 3 | Output | Precentral gyrus | 48 | 0 | 45 | 0.538 | 59 | -0.405<sup>*</sup> | -0.178 | -0.136 | -0.075 | -0.261 |
| Amygdala | Input | Posterior insula | 39 | -9 | 18 | -0.507 | 27 | 0.414<sup>*</sup> | 0.241 | 0.048 | 0.047 | 0.205 |
| ***P-PTS*** |  |  |  |  |  |  |  |  |  |  |  |  |
| Anterior cingulate cortex | Output | Middle temporal gyrus | -45 | -75 | 15 | 0.447 | 34 | -0.256 | -0.319 | 0.032 | -0.107 | -0.085 |
| Dorsomedial prefrontal cortex 1 | Output | Precentral gyrus | 45 | 0 | 45 | 0.514 | 46 | -0.239 | -0.229 | -0.001 | 0.100 | -0.121 |
| Dorsomedial prefrontal cortex 2 | Input | Posterior cingulate cortex | -3 | -45 | 18 | 0.453 | 34 | -0.160 | -0.190 | 0.173 | 0.030 | 0.023 |
| Dorsomedial prefrontal cortex 3 | Input | Posterior cingulate cortex | 3 | -45 | 15 | 0.547 | 126 | -0.258 | -0.075 | 0.050 | -0.049 | -0.152 |
| Dorsomedial prefrontal cortex 3 | Output | Inferior parietal lobe | 39 | -42 | 57 | 0.496 | 37 | -0.188 | -0.044 | -0.091 | -0.233 | -0.179 |
| Dorsomedial prefrontal cortex 3 | Output | Precentral gyrus | 48 | 0 | 45 | 0.583 | 84 | -0.375 | -0.221 | -0.153 | -0.055 | -0.262 |
| Amygdala | Input | Posterior insula | 39 | -9 | 21 | -0.542 | 78 | 0.451<sup>*</sup> | 0.225 | 0.036 | 0.090 | 0.230 |
| Amygdala | Input | Supramarginal gyrus | 60 | -42 | 39 | 0.498 | 27 | -0.395<sup>*</sup> | -0.137 | 0.131 | -0.149 | -0.163 |

a: Number of voxels. FWE ( $p < 0.05$ ) with voxel- $p < 0.001$ . Voxel size = $3 \times 3 \times 3$ .
T-PTS: total psychopathic traits score; P-PTS: primary psychopathic traits score.
\*: Bonferroni-corrected $p < 0.05$ , $n = 65$ .

### Relationship between concern with moral foundations and EC that were significantly related to psychopathic traits

Partial correlation analyses were further conducted between the EC that was significantly associated with psychopathic traits and concern with the Harm foundations. For the EC that was significantly correlated with T-PTS, a positive correlation between the EC from the posterior insula to the amygdala and the Harm foundation and a negative correlation between the EC from the dmPFC to the precentral gyrus and the Harm foundation was found. For the EC that was significantly correlated with P-PTS, a positive correlation between the EC from the posterior insula to the amygdala and the Harm foundation, and a negative correlation between the EC from the amygdala to the supramarginal gyrus and the Harm foundation were found. However, no correlations were found between the other four foundations and the EC that was significantly related to psychopathic traits (see **Tab. 4**).

### Mediation analysis

For EC significantly correlated with both psychopathic traits and concern with the Harm foundation (i.e., EC from the posterior insula to the amygdala, and from the dmPFC to the precentral gyrus), mediation analyses were further performed to investigate whether the EC mediated the relationship between psychopathic traits and the concern with the Harm foundation. The indirect effect of T-PTS through the EC from the posterior insula to the amygdala (95% CI = [-0.150, -0.003]) on the Harm foundation was significant. The direct effect of T-PTS on the Harm foundation was still significant when the EC was included as a mediator (95%CI = [-0.268, -0.025]), indicating that the EC from the posterior insula to the amygdala partially mediated the relationship between T-PTS and concern with the Harm foundation. No mediation effect of T-PTS through the EC from the dmPFC to the precentral gyrus on the Harm foundation was observed (95%CI = [-0.128, 0.169]) (**Fig.4**).

**Figure 4.**
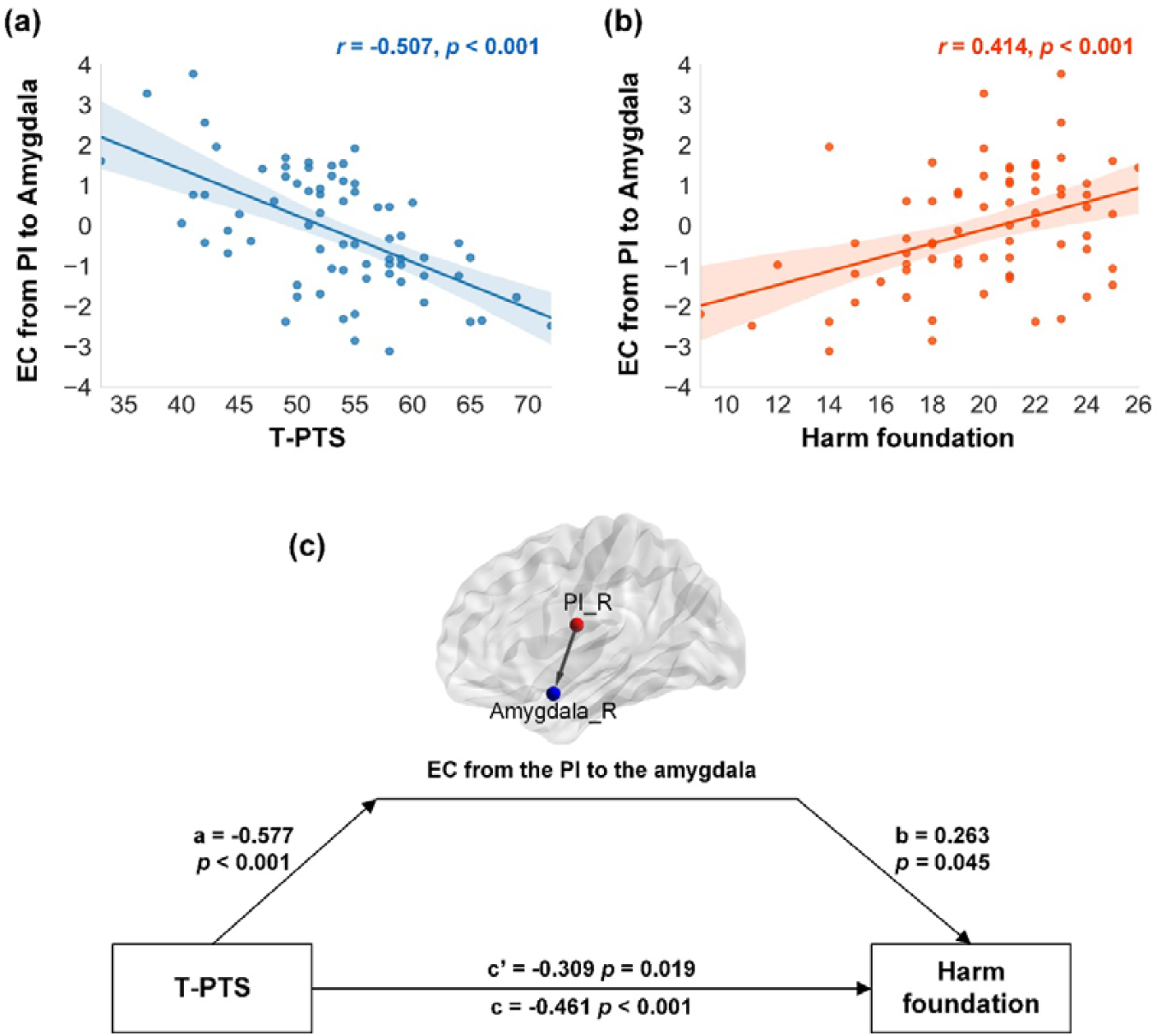
Correlation and mediation analyses (T-PTS). (a) EC from the insula to the amygdala was negatively correlated with T-PTS. (b) EC from the posterior insula to the amygdala was positively associated with moral concern with the Harm foundation. (c) EC from the insula to the amygdala partially mediated the association between T-PTS and moral concern with the Harm foundation. EC: effective connectivity; PI: posterior insula; PI_R: right posterior insula; Amygdala_R: right amygdala; T-PTS: total psychopathic traits score.

Similarly, the indirect effect of P-PTS through the EC from the posterior insula to amygdala (95%CI = [-0.400, -0.041]) on the Harm foundation was significant, but the indirect effect of P-PTS through EC from the amygdala to the supramarginal gyrus (95%CI = [-0.163, 0.020]) was not significant, Moreover, the direct effect of P-PTS on the Harm foundation was still significant when the EC was included as a mediator (95%CI = [-0.249, -0.001]), indicating that the EC from the posterior insula to the amygdala partially mediated the relationship between P-PTS and concern with the Harm foundation (**Fig. 5**).

**Figure 5.**
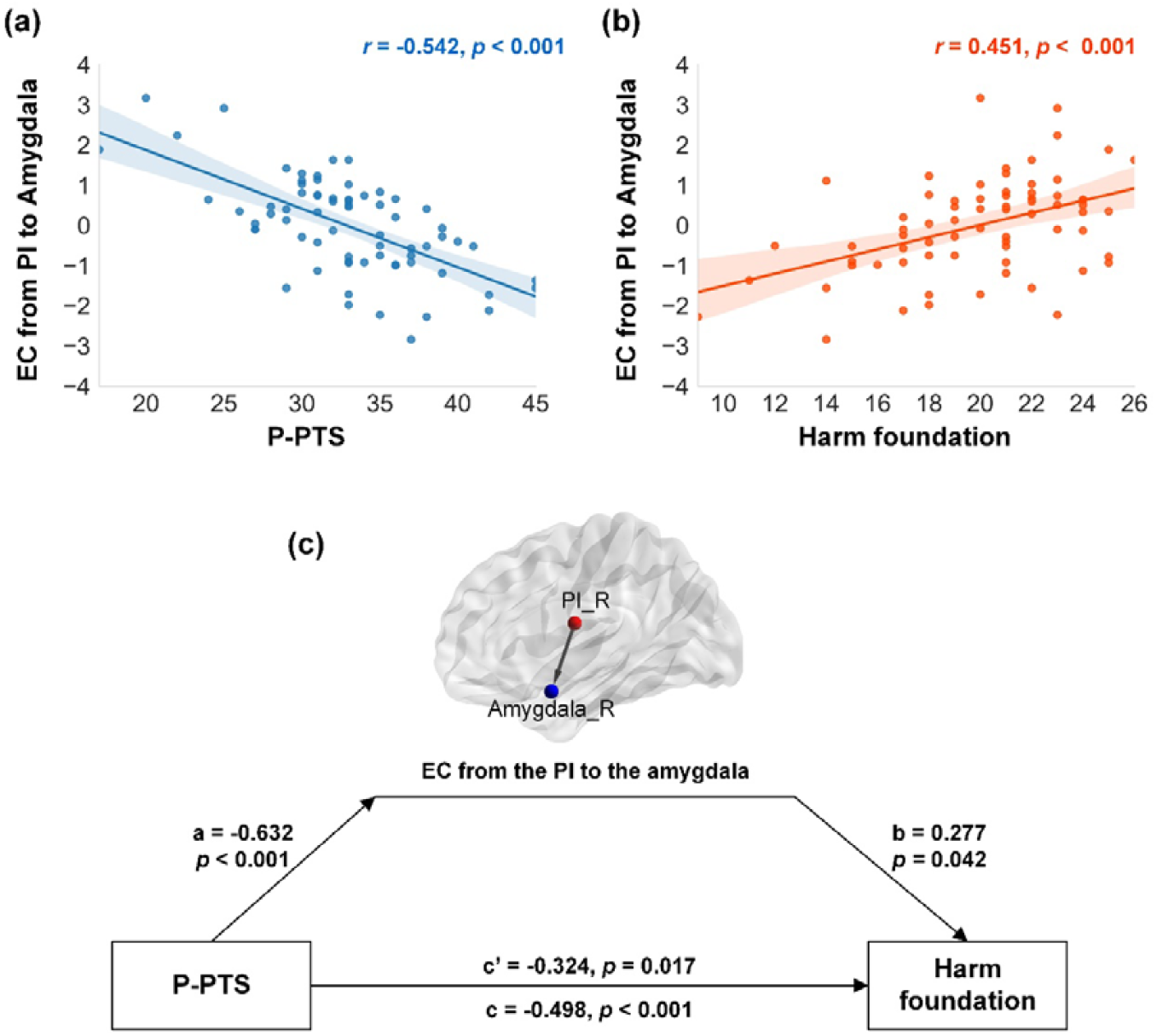
Correlation and mediation analyses (P-PTS). (a) EC from the insula to the amygdala was negatively correlated with P-PTS. (b) EC from the insula to the amygdala was positively associated with concern with the Harm foundation. (c) EC from the insula to the amygdala partially mediated the association between P-PTS and concern with the Harm foundation. EC: effective connectivity; PI: posterior insula; PI_R: right posterior insula; Amygdala_R: right amygdala; P-PTS: primary psychopathic traits score.

## DISCUSSION

This study combined self-report questionnaires with RS-fMRI EC to reveal the intrinsic neuropsychological mechanism of the relationship between psychopathic traits and concern with different moral foundations. At the behavior level, we found that T-PTS and P-PTS predicted moral concern with the Harm foundation, but not the other four foundations. At the neural level, we found that reduced EC from the posterior insula to the amygdala was associated with higher psychopathic traits and greater concern for the harm foundation. Furthermore, the EC from the posterior insula to the amygdala partially mediated the relationship between T-PTS/P-PTS and concern with the Harm foundation. Our findings provided the first neural evidence for less concern with the harm foundation for individuals with higher psychopathic traits and deepened our understanding of the underlying mechanism of atypical moral cognition associated with psychopathic traits.

Our behavioral results demonstrated that T-PTS and P-PTS significantly predicted moral concern with Harm foundation only. This is consistent with previous studies which reported significant association between psychopathic traits and different moral foundations, with the effects being most pronounced for the Harm foundation (Aharoni et al., 2011; Gay et al., 2018; Glenn et al., 2009a; Marshall et al., 2018). Perception of harm is intuitive, forming the fundamental basis of moral judgment (Haidt & Joseph, 2004; Schein & Gray, 2017). A systematic review suggested that psychopathy is primarily associated with compromised care-based morality involving moral reasoning about harmful actions to others (Blair, 2007). Here, we provided more evidence that psychopathic traits are more related to harm-based moral cognition than other types of moral cognition.

Our neural results showed that decreased EC from the posterior insula to the amygdala was associated with higher psychopathic traits. The amygdala and insula are core components of the salience network (SN), a large-scale brain network widely recognized to be involved in detection and integration of emotional and sensory stimuli (Geng et al., 2016). Both of the two regions take a crucial role in saliency processing, attention capturing and identification of emotional significance of the stimulus (Calder et al., 2001; Menon & Uddin, 2010; Phillips et al., 2003). Since dysfunctional emotional responses are characteristic features of individuals with higher psychopathic traits, atypical neural activity in the amygdala and insula is very likely to be associated with psychopathic traits (Blair, 2013; Dolan & Fullam, 2009; Ermer et al., 2012; Santana, 2016; Yang et al., 2009). For example, a systematic review suggested that brain regions implicated in psychopathic traits include the amygdala, insula, anterior and posterior cingulate, etc. (Santana, 2016). Individuals with higher psychopathic traits show reduced amygdala and insula volume (Ermer et al., 2012; Yang et al., 2009), as well as decreased neural activity in the amygdala and insula (Blair, 2013; Dolan & Fullam, 2009). Furthermore, individuals with higher psychopathic traits show altered anatomical and functional connectivity between the two regions and other brain areas, and between the two regions themselves during socio-moral decision making (Fumagalli & Priori, 2012; Han et al., 2016; Shenhav & Greene, 2014).

The insula is anatomically and functionally connected with the amygdala (Augustine, 1985, 1996; Gasquoine, 2014; Sethi et al., 2018; J. L. Stein et al., 2007). This pathway constitutes a part of the network associated with salience and emotion processing (Baur et al., 2013; Geng et al., 2016). The insula has been commonly recognized to translate bodily state information into emotional feelings (Dennis et al., 2011; Uddin, 2014; Yu et al., 2015). The interoceptive signals arrived in the posterior insula before being evaluated and represented in the anterior insula (Craig & Craig, 2009; Straube & Miltner, 2011). The right posterior insula is considered to be especially involved in the awareness of one’ own bodily states. When people pay more attention to their own emotions, the activities of the primary somatosensory cortex as well as the posterior insula increase (Straube & Miltner, 2011). The amygdala receives inputs of various sensory and emotional information from other brain regions (Diano et al., 2017; Hofmann & Straube, 2019) including the posterior insula. Evidence showed that resting-state connectivity between the amygdala and the posterior insula decreased with emotional dysregulation (Bebkoa et al., 2015). The amygdala may detect and integrate the bodily state information transmitted from the posterior insula to support adaptive emotional responses (Babaev et al., 2018; Bebkoa et al., 2015; Stein et al., 2007). We argued that as the EC from the posterior insula to the amygdala decreases, the capacity to recognize and integrate bodily state information into emotional responses might weaken for individuals with higher psychopathic traits.

Furthermore, we found that the EC from the posterior insula to the amygdala partially mediated the relationship between psychopathic traits and the concern with the Harm foundation. In conjunction with previous behavioral findings that emotion responses serve as a mediator in the relationship between psychopathic traits and morality (Glenn et al., 2009; Patil, 2015; Ye et al., 2021), the results here suggested that weakened capacity to integrate bodily-state information to emotional responses in individuals with elevated psychopathic traits in response to moral contexts accounts for their emotional shallowness, which partially contributes their atypical moral cognition. Importantly, the significant relationship between psychopathic traits and morality as well as the mediating role of the posterior insula-to-amygdala connectivity in the relationship was only observed in T-PTS and P-PTS, but not S-PTS. These results concur with previous findings that only primary psychopathy is able to predict moral judgments in the contexts of sacrificial moral dilemmas (Takamatsu & Takai, 2019) and every-day moral scenarios (Ye et al., 2021). Primary psychopathy covers interpersonal and affective features, whereas secondary psychopathy represents aggressive and antisocial lifestyles (Hare, 1991). The primary psychopathic subtype normally demonstrates significantly lower anxiety than the secondary subtype (Lee & Salekin, 2010; Skeem et al., 2007; Vaillancourt & Brittain, 2019). In addition, reduced amygdala and insula activity in response to fear stimuli is merely associated with the primary psychopathy (Sethi et al., 2018). Overall, our findings supported that primary psychopathy contributes remarkably more than the secondary psychopathy to the atypical moral cognition and the neural underpinnings of the relationship of the two subtypes of psychopathic traits to moral cognition might differ significantly.

It is noteworthy that the aforementioned neural finding (i.e., the posterior insula-amygdala EC) was restricted to the right hemisphere which is more frequently involved in the processing of all kinds of emotions, especially negative emotions (Bowu et al., 1988; Gainotti, 2019; Killgore & Yurgelun-Todd, 2007). For example, increased activations were observed in emotion-related brain regions anchored in the right hemisphere (e.g., the right vmPFC and right amygdala) during a facial emotion perception task, especially in the case of facial cues with negative emotions (Killgore & Yurgelun-Todd, 2007). Thus, the EC from the posterior insula to the amygdala related to psychopathic traits and morality was only found in the right hemisphere may be due to the lateralized processing of negative emotion.

Additionally, we found that the EC from the dmPFC to the precentral gyrus was significantly correlated with T-PTS and P-PTS. Increased EC from the dmPFC to the precentral gyrus was associated with higher psychopathic traits. The dmPFC contributes to cognitive control and goal-oriented action selection (Ridderinkhof et al., 2004; Venkatraman et al., 2009) whereas the precentral gyrus is responsible for the control of voluntary motor movement (Loukas et al., 2011). The altered EC from the dmPFC to the precentral gyrus might be linked to poorer behavioral control of individuals with higher psychopathic traits compared to those with lower psychopathic traits who are more capable of appropriately taking goal-oriented actions. However, we did not find any mediating effect of the EC from the dmPFC to the precentral gyrus on the relationship between psychopathic traits and morality. Therefore, the increased communication from the dmPFC to the precentral gyrus might be part of the intrinsic neural characteristics for psychopathic traits but not the potential neural mechanism for how psychopathic traits were linked to morality.

Several limitations of this study are noteworthy. First, only college students were included as the participants and the sample size was relatively modest. Replication in a larger community sample is needed in the future to enhance the generalizability of our findings. Second, some items in the MFQ items seem rather abstract and obscure for participants, leading to relatively low validity and reliability of the scale. More accessible measures for morality (e.g., daily moral scenarios) should be utilized in the future to improve the reliability and especially the external validity of the measurement (Clifford et al., 2015; Ye et al., 2021). Third, ROIs used in the study were from a meta-analysis, which may bias the results as we may overlook other important brain areas (vmPFC, posterior cingulate, and superior temporal gyrus) that are related to psychopathic traits. In addition, the repetition time of RS-fMRI in our study was 2 s which is much slower than the neuron reaction time. However, many researchers believe that even in the case of using long TRs to collect fMRI data, GCA still estimates the time-directional influences (Liao et al., 2010; Rajan et al., 2019). Finally, future studies should also examine the relationship between psychopathic traits and moral concern in the Liberty foundation which was recently validated as a new moral foundation (Graham et al., 2018).

Taken together, the present study revealed the underlying neuropsychological signatures of the relationship between psychopathic traits and concern with different moral foundations. Our findings demonstrated that T-PTS and P-PTS predicted concern with the Harm foundation. Importantly, higher psychopathic traits were associated with reduced EC from the posterior insula to the amygdala, which partially mediated the relationship between psychopathic traits and concern with the Harm foundation. These findings provided new evidence for the special role of psychopathic traits in concern with different foundations and shed new light on the neuropsychological mechanisms underlying the relationship between psychopathic traits and morality. This study increases our knowledge about the atypical neural circuits associated with moral cognition for higher-psychopathic-trait individuals in terms of effective connectivity between brain regions and may have some implications for early prevention of individuals with higher psychopathic traits from committing serious transgressions.

